# Sub-operon promoter arrangement of *disA* facilitates c-di-AMP homeostasis and selective stress responses in *M. smegmatis*

**DOI:** 10.1101/2022.06.29.498085

**Authors:** Mamta Singla, Aditya Kumar Pal, Vikas Chaudhary, Anirban Ghosh

## Abstract

Bacterial second messenger signaling often plays an important role in cellular physiology. In this study, we have attempted to understand how c-di-AMP synthesis and degradation are transcriptionally regulated in *M. smegmatis*. We have discovered that, although c-di-AMP synthesis gene *disA* exists in a multi-gene operon; a sub-operon promoter arrangement plays a key role under various stress conditions, keeping its dual function property intact. Further, we learned that c-di-AMP plays a role in the autoregulation of the *disA* promoter to limit intracellular c-di-AMP concentration. We also identified an alternate start codon within the *disA* gene which can lead to the synthesis of truncated DisA protein at times using an independent stress-inducible promoter. All in all, this study was helpful to understand how c-di-AMP synthesis is regulated under normal and stress conditions linked to its physiological relevance in *M. smegmatis*.

## Introduction

Cyclic-di-adenosine monophosphate (c-di-AMP) is an essential second messenger in many Gram-positive bacteria known to coordinate important cellular functions (Corrigan et al., 2011; Corrigan et al., 2013; Luo and Helmann, 2012; Oppenheimer-Shaanan et al., 2011). In Mycobacteria, the c-di-AMP level is constitutively maintained across all growth phases at a minimum concentration (Tang et al., 2015), but under relevant stress conditions, intracellular c-di-AMP levels could be modulated by the upregulation of the *disA* gene, which encodes for the synthetase enzyme to convert 2 ATP molecules into c-di-AMP. It has been previously reported that the *disA* gene in *M. smegmatis* exists in a multi-gene operon along with two other genes *radA* and *lpqE* (Zhang and He, 2013). Whereas the *lpqE* is a gene so far functionally uncharacterized, DisA and RadA proteins have been studied well and found to have a definite role in DNA damage response when cells encounter some genotoxic stress (Gandara and Alonso, 2015; Torres et al., 2019; Witte et al., 2008). Apart from being the sole DAC (Diadenylate cyclase) enzyme responsible for c-di-AMP synthesis, DisA functions as a DNA integrity scanning enzyme, which might have an undisclosed role in DNA repair and mutagenesis(Bejerano-Sagie et al., 2006). Based on a previous study, RadA and DisA are thought to be co-transcribed using the common operon promoter present upstream of the operon (Zhang and He, 2013). The same study claims that RadA protein inhibits the synthetic activity of DisA by physically interacting with the DisA enzyme (Zhang and He, 2013) and possibly directing its function exclusively to DNA damage repair, further emphasizing the fact that these two proteins are functionally linked. On the other hand, a low yet constant level of c-di-AMP synthesis and maintenance is important for cellular physiology (Mehne et al., 2013), which is only feasible if there is an alternative mechanism to ensure that the c-di-AMP synthetic function of the DisA enzyme remains uninhibited. One possibility of that scenario will be when the *disA* gene can be expressed independently of *radA* using an alternate promoter just upstream of its own ORF. Fresh Bioinformatics and a recent study have revealed the presence of a potential transcription start site (TSS) 25 bp upstream of the *disA* start codon (Martini et al., 2019). This observation makes us curious if there is any sub-operon arrangement exists (Li et al., 2017) and whether the independent promoter element which is present just upstream of the *disA* gene can be active during normal growth and if at all inducible under specific stress conditions where c-di-AMP plays a major role. Apart from that, we were also curious to study the autoregulation, possible translational repression of the *disA* promoter and if the different domains of this complex enzyme DisA prefer to have independent expression and activity relevant to some stresses.

## Materials and Methods

### Bacterial strains, media and growth conditions

All the bacterial strains used in the study are listed in **Table S1**. *E. coli* strains were grown in Lysogeny broth (LB) media. *M. smegmatis* mc^2^ 155 and its derivatives were grown at 37°C in Middlebrook 7H9 (MB7H9) media (Himedia) supplemented with 2% (v/v) glucose, 0.05%(v/v) tween, and 50 µg/ml of Hygromycin (SRL) unless stated otherwise (Sharma et al., 2014). For agar plates, 1.5% agar was added to the liquid medium (without tween).

### Construction of Transcriptional Fusion Constructs

All the reporter constructs were made in promoter-less GFP vector pMN406-ΔPimyc (Roy et al., 2004) (a generous gift from Prof. Ajitkumar, Indian Institute of Science, Bengaluru). To generate transcriptional fusions, different fragments from the upstream regions of the operon genes *MSMEG_6078 (lpqE), MSMEG_6079 (radA), MSMEG_6080 (disA)* as well as truncated *disA* were PCR amplified from *M. smegmatis* mc^2^155 genomic DNA using a compatible set of primers. These fragments were then cloned into the upstream of the GFP in the reporter vector using different restriction enzymes. The generated plasmids were verified by the sequencing and were electroporated into *M. smegmatis* mc^2^155 cells, and the transformants were selected against hygromycin (50 µg/ml). All the restriction enzymes used in cloning were purchased from New England Biolabs (NEB). All the primers used to make plasmid constructs are listed in **Table S2**.

### Measurement of Growth and Base promoter activity

As mentioned by (Scholz et al., 2000), to measure the GFP induction in the various transformants, 200 µl of cells were taken in a black, flat bottom 96 well plate (Thermo Nunc) and GFP intensity was recorded at excitation of 488 nm and emission of 510 nm in Varioskan Flash multimode reader (Thermo Fisher Scientific) bandwidth was set to 5nm. GFP expression in Relative Fluorescence Unit (RFU) was calculated by dividing fluorescence intensity Fluorescence Unit (FU) by cell density (OD_600_). *M. smegmatis* harboring pMN406-ΔPimyc vector was used as negative control and *M. smegmatis* (pMN406-ΔPimyc-*hsp60*) was used as the positive control where required. All the measurements were corrected against the blank and were performed in three biological replicates. To characterize the putative promoter region for *msdisA*, the cells without any exposure to stress were used as the reference for the induction profile. For this, the cells were grown till the mid-late exponential phase, further inoculation was done in standard growth conditions with 2% glucose to make the final OD_600_ nm of ∼0.03. The bacterial growth and GFP intensity were further monitored (as described in the material and methods) at different intervals for 84 -108 hours.

### Promoter Induction estimation during stress

For the Glucose and pH adaptation experiments, cells from the mid-late exponential phase were harvested and washed twice in fresh MB7H9 media containing 0.05% tween 80. Subsequently, the pellet was resuspended in media containing different concentrations of glucose from the standard (2%) to starved (0.2%) conditions, or the MB7H9 media with acidic pH 4 or basic pH 9 and adjusted to OD_600_ ∼0.03. For stress response experiments, cells were harvested at log phase OD_600_ 0.6-0.8 and then exposed to a different category of stresses and kept at a 37°C shaker incubator post-treatment. Individual promoter induction was measured and quantitated at subsequent timepoints. Ciprofloxacin and KCl were used at final concentrations of 1µg/ml and 500 mM respectively, whereas, in case of UV stress, cells were exposed to a UV dose of 0.15 mJ/cm^2^. The fluorescence was measured every 4 hours up to 24 hours to check the stress-induced promoter induction in a kinetic manner.

### Stress survival assays

*M. smegmatis* strains were grown in presence of appropriate antibiotics until the mid-log phase and then normalized to OD_600_ 0.7. 1 ml. of this culture was exposed to UV irradiation (0.15 mJ/cm^2^) and kept at a 37°C shaker incubator for 4 hours. The bacteria were then diluted by 10-fold serial dilutions and 4 μl of cells from each dilution were spotted into MB7H9 agar medium, incubated at 37°C for 3 days and CFU/ml. was calculated. For NaCl treatment, similar dilution series were made from OD_600_ 0.7 culture of the respective strains and 20 μl of culture from selective dilutions were spread on MB7H9 agar plates containing 500 mM NaCl. Survival percentage was calculated as the percentage of CFU remaining after exposure to respective stresses compared to the CFU in the untreated control. Antibiotic stress was estimated by the disc diffusion assay. Briefly, *M. smegmatis* strains were grown until the mid-log phase and 100 μl of the OD_600_ 0.7 cultures were spread on MB 7H9 agar plates and left for drying. After that, ciprofloxacin discs (2.5 mg/ml.) were aseptically kept in the middle of the plate using a sterile tweezer. Plates were further incubated at 37°C for 3 days and the zone of inhibition (cm^2^) was calculated as a measurement of antibiotic sensitivity.

### Microscopy

For microscopic analysis, cells were exposed to UV irradiation and KCl treatment (Dose and concentration mentioned before) and were harvested at 0- and 4 hours post-treatment, washed with 1X PBS (pH 7.4) twice, and concentrated. It was followed by fixation with 1% paraformaldehyde for 4 hours in dark [14]. Fixed cells were then placed on a glass slide and observed under a confocal microscope (Leica TCS SP8, Hyvolution). Images were processed using the LASX software.

## Results

### The *disA* operon structure and identification of gene-specific sigma factor recognition sites, transcription start sites and RBS

It has been shown in one of the previous studies that *radA* and *disA* gene exists in the same operon and are often co-transcribed (Zhang and He, 2013), but based on the DOOR3 database (Cao et al., 2019), operon prediction for *M. smegmatis* mc^2^155 (NCBI Genbank accession NC_008596) *disA* gene was not predicted to be in an operon. Therefore, we decided to have a relook of the entire operon sequence for any potential promoters that exist inside the operon and/or specific to each gene based on the mycobacterial sigma factor promoter consensus recognition sites (Newton-Foot and Gey van Pittius, 2013). By using the tool genome_pattern_search (http://www.ualberta.ca/~stothard) and manual inspection we identified several potential *sigB* and *sigE* consensus recognition sites within the intergenic regions of *lpqE-radA-disA* genes, which could theoretically act as RNA polymerase attachment sites to facilitate transcription of downstream genes (Newton-Foot and Gey van Pittius, 2013) **(Fig.1a)**. After this, based on a RNA-seq data in a previous publication (Martini et al., 2019) we identified the independent transcription start sites for all 3 genes and it was assumed that the RNA polymerase could potentially initiate the transcription of individual genes without depending on the upstream gene’s TSS. Next, we checked the possible shine-Dalgarno sequence (RBS) ahead of the start codons of all 3 genes in the operon and found perfect RBS sequences upstream of *lpqE* and *disA*, but not *radA* **(Fig.1b)**. These critical observations could imply that at times these genes could be expressed independently of each other, which led us to investigate further this sub-operon promoter arrangement of the *lpqE-radA-disA* operon.

**Figure 1.**
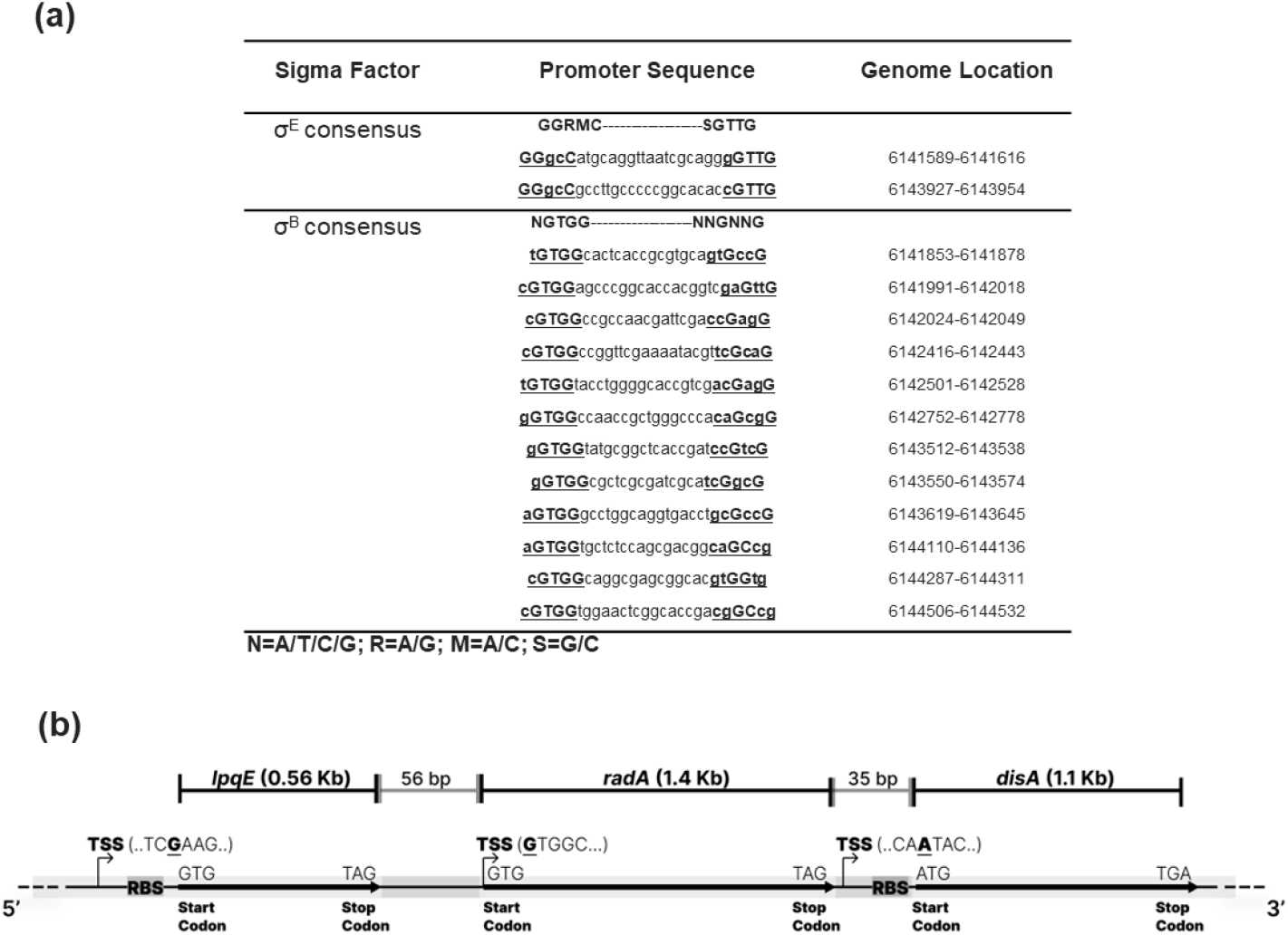
**(a)** Putative sigma factor sites in inter/intergenic regions are presented in the table. The σ^E^ and σ^B^ sequences used for the search are shown on the first line. The consensus sequence calculated from the mycobacterial sequences illustrated is shown below the alignments. Putative −10 and −35 sequences are underlined, and bases matching the mycobacterial consensus sequence are in bold. Bases of the consensus sequence conserved in all sequences are in capital letters. The numbers on the right refer to the position of the consensus sequences in *M. smegmatis* MC^2^155 genome. **(b)** Genetic organization of *disA* operon in *M. smegmatis*; Predicted transcription start sites (TSS), ribosome binding sites (RBS) and start codons are highlighted for *lpqE, radA* and *disA* genes.

### Studying the Core and extended promoter region of *disA* gene

As per our bioinformatic analysis of *disA*’s independent promoter, we cloned the 300 bp upstream region of the *disA* gene into a promoter-less GFP vector to make a transcriptional fusion construct (Roy et al., 2004). To have an unbiased and specific outcome, we also made two similar transcriptional fusion constructs by taking either 1 Kb or 2.3 Kb upstream (which comprises the entire operon sequence) DNA region of the *disA* gene in the mycobacterial chromosome **(Fig.2a)**. We grew the cells harboring the three different promoter-fusion constructs in standard in MB7H9+tween+glucose and measured the OD_600_ and GFP intensity at different growth phases. Though there was understandably no difference in cell growth (OD_600_), it was observed that the 300bp-us region of *disA* had the highest promoter activity (GFP intensity) followed by 1 kb and 2.3 kb full-length construct **(Fig.2b)**. P_*hsp60*_-GFP construct has been used as a positive control (Dellagostin et al., 1995) and promoter-less empty vector served as a negative control in the assay. Further, we checked the individual promoter activity under different stress conditions relevant to c-di-AMP’s known physiological roles in different bacteria (Commichau et al., 2015; Yin et al., 2020). Specifically, kinetic measurements of OD_600_ and GFP intensity were taken up till the late stationary phase and it was observed that when the media was supplemented with 10 times less glucose of 0.2% f.c. (carbon starvation), 300 bp us region showed significant induction compared to others, particularly in the stationary phase **(Fig.2c)**. As expected, cell growth (OD_600_) was seen to be compromised under low glucose conditions and the timing of the growth stasis and promoter induction was having a high correlation. Next, we tried to check the promoter induction if the cells are grown in a media with non-optimal pH (acidic pH 4.0 and basic pH 9.0) and found that 300us *disA* promoter induction increased by 2-3-fold during growth in acidic pH 4 media **(Fig. 2d)**. At pH 9, a modest increase was observed compared to the pH 7 control series, but it seems non-significant (data not shown). Thus, our consolidated data of the *disA* core (300bp) promoter induction pattern suggested that the *disA* gene can be transcribed using an alternate transcription start site under normal physiological conditions to maintain the critical concentration of c-di-AMP inside cells as well as under specific stress conditions, which could be linked to the corresponding stress adaptation. Comparative GFP expression data suggested that the extended promoter regions of *disA* do not get induced significantly under the same stresses and might have some regulatory components to prevent *disA* expression when the whole fragment is transcribed using any upstream TSS **(Fig. S1)**.

**Figure 2.**
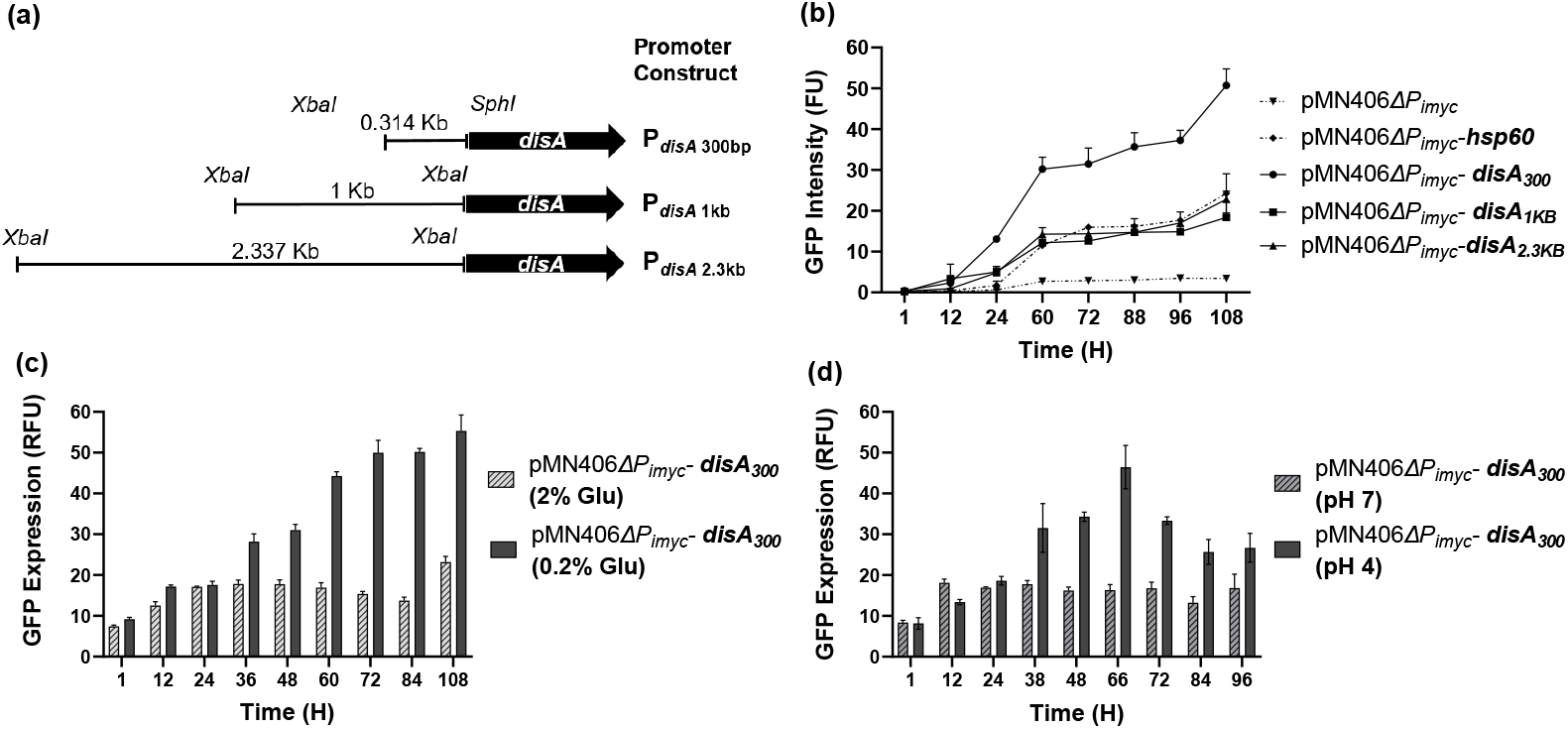
**(a)** Schematic representation of different *disA* promoter fusion constructs along with length mentioned on top. **(b)** Promoter activity of different *disA* promoter fusion constructs are measured under normal conditions of growth as a function of GFP fluorescence over time. Promoter induction of the P_*disA* 300 bp_ construct is measured in kinetic manner upon carbon stress **(c)** and acidic pH 4 **(d)** upto 108 hours. The graphs were plotted using GraphPad Prism 9. *** = P < 0.001; ** = P < 0.01; * = P < 0.05.

### Stress-specific induction of core promoter regions of *radA* and *disA* genes

As our bioinformatics analysis and previous publication (Martini et al., 2019) suggested the presence of distinct TSS and sigma factor recognition sequences upstream of all three genes in the operon, we were interested to study how these individual genes’ core promoter regions (∼300-400 bp us of the corresponding start codons) respond to different stress conditions. Since our previous observation has hinted at a considerable expression of GFP only from the core promoter region (300bp us) of *disA*, we have taken the core promoter regions of *radA* and *lpqE* genes to make similar GFP-based transcriptional fusion constructs. These three constructs: 400us-*lpqE*, 300us-*radA* and 300us-*disA*, also referred to as P1, P2, and P3 **(Fig. 3a)** were designed to study selective inductions under specific stress conditions, which are physiologically relevant to either DisA enzyme function or c-di-AMP signaling pathway. Based on a previous report of c-di-AMP’s role with UV irradiation in *M. smegmatis* (Manikandan et al., 2018), we treated the cells harboring different reporter constructs to genotoxic stresses such as UV irradiation (0.15 mJ/cm^2^) and measured the corresponding GFP fluorescence after 4 hours (∼1 generation time) of incubation at 37°C. Previously, we found that *ΔdisA* strain (c-di-AMP null strain) survives better under ciprofloxacin treatment by some unidentified mechanism (unpublished data). Relating both the contexts, we observed significant induction of P2 (P_*radA*_) promoter elements under DNA-damaging conditions, which were previously not identified. Compared to untreated control there was a 3-4 fold increase in induction of GFP fluorescence indicating moderately strong promoter induction when cells were exposed to UV and ciprofloxacin respectively **(Fig. 3b & 3c)**. When cells were further incubated, the induction profile remains unchanged for up to 24 hours **(Fig. S2a & S2b)**. In the case of the *disA* core promoter (P3), a moderate induction was observed in the case of UV stress and no induction was observed with ciprofloxacin treatment, implying the selective induction of the *disA* gene under DNA damaging conditions was not favoured by the cell **(Fig. 3b & 3c)**. Surprisingly, there was no significant promoter induction observed in the case of P1 (*lpqE* core promoter) **(Fig. 3b & 3c)**, which was considered to be the upstream operon promoter. Selective induction of only the *radA* core promoter (P2) highlighted the role of *radA* gene and possibly downstream gene *disA* (between them there is no transcription termination sequence) as a part of the DNA repair response network. Next, we checked the increase of GFP fluorescence at the single-cell level to reveal if there was any cellular heterogeneity involved in the promoter induction. Our microscopy data suggested that there was a significant difference in induction of the r*adA* core promoter after 0.15 mJ/cm^2^ UV exposure compared to the untreated control **(Fig. S3)**. UV exposure also caused a visible fluorescence increase in the case of *disA* promoter **(Fig. S3)**, further corroborating our previous observation at the population level **(Fig. 3b)**. In the case of both strains, after UV exposure, elongated cells were predominantly visualized because of SOS DNA repair-driven cell cycle arrest, as part of the global response to DNA damage (Janion, 2008).

**Figure 3.**
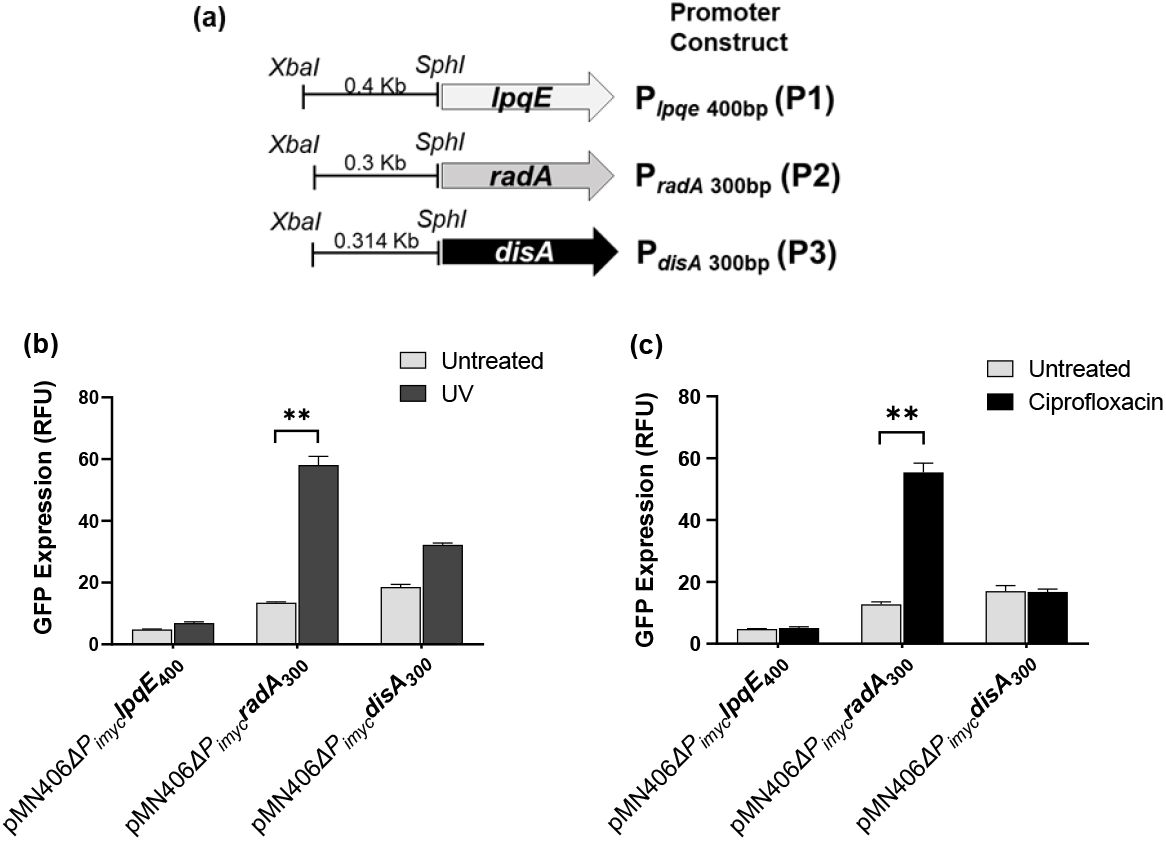
**(a)** Schematic representation of different core promoter fusion constructs of *lpqE, radA* and *disA* genes, along with the gene names and length mentioned on top. Individual promoter activity of P_*lpqE* 400bp_, P_*radA* 300bp_ and P_*disA* 300bp_ constructs are measured upon **(b)** 0.15 mJ/cm^2^ UV irradiation and **(c)** 1μg/ml Ciprofloxacin treatment, readings are taken 4 hours post exposure. The graphs were plotted using GraphPad Prism 9. *** = P < 0.001; ** = P < 0.01; * = P < 0.05.

Next, we checked the individual constructs’ promoter induction when cells were exposed to ionic stress in liquid broth. Cells were grown till the mid-logarithmic phase of OD∼0.7 and KCl and NaCl were added to the final conc. of 500 mM and GFP intensity was checked after 4 hours of incubation at 37°C. In the case of P1 and P2 (*lpqE* and *radA*) constructs, no increase in GFP fluorescence was observed compared to untreated control over 24 hours. Though we have observed a small increase in P3 (*P*_*disA*_) promoter activity in the case when cells were treated with 500 mM KCl after 4 hours compared to the untreated control, the data were not reproducible after multiple trials **(Fig. S4a)** and hence not considered significant. When we checked the induction with a more sensitive fluorescence microscopy imaging, we could not find a visible difference in fluorescence intensity either, between untreated and KCl (500 mM) treated samples after 4 hours of treatment **(Fig. S4b)**. To understand, the specific dose-dependency of salts, we also checked the P3 induction at a lower salt concentration of 250 mM and found the promoter remained uninduced (data not shown). To our surprise, the *radA*(P2) promoter, which showed significant induction under multiple stresses before, remained unchanged with both 250 mM and 500 mM KCl treatment **(Fig. S4a)** and microscopic images also confirmed no increase in GFP and change in cell morphology. As we know, increased c-di-AMP concentration is detrimental for cells when they encounter salt stress (Huynh and Woodward, 2016), conversely, we checked the induction profile of P_*pde*_ to find out if more Pde protein (hydrolyses c-di-AMP) copy number and enhanced degradation of c-di-AMP could be the controlling factor responsible for salt stress tolerance. But our data suggested that there was no significant induction or repression of P_*pde*_ under salt stress either **(Fig. S4c)**, further suggesting that c-di-AMP synthesis and degradation genes are not transcriptionally regulated upon ionic stress in *M. smegmatis*.

### Overexpression of RadA does not result in better survival of the strain signifying its regulatory role

Based on a previous study (Manikandan et al., 2018), it was already known that in *M. smegmatis*, c-di-AMP binds to RecA protein and attenuates the nucleoprotein complex formation resulting in a defective SOS response upon DNA damage. So, when a bacterial cell encounters DNA damage induced by UV irradiation or ciprofloxacin (Cordone et al., 2011; O’Sullivan et al., 2008) they prefer to repair the breaks through a potentially error-free RecA-dependent homology-based repair system to avoid unnecessary mutation (Heidenreich et al., 2003). In such conditions, from a cell’s point of view, it becomes a priority to keep the intracellular concentration of c-di-AMP down by inhibiting DisA’s synthesis function. Selective induction of the *radA* core promoter, but not the downstream *disA* core promoter under such conditions is critical to elevate the RadA enzyme concentration to sufficient levels to ensure inhibition of c-di-AMP biosynthesis by DisA. To find out whether a higher copy number of RadA plays any significant role in DNA repair by itself, the *radA* gene was overexpressed from a multi-copy plasmid pMV261 (Stover et al., 1991) and data suggested no better survival compared to WT with pMV261-empty plasmid upon UV exposure **(Fig. 4a)**. Similarly, when we overexpressed the *disA* gene from the pMV261 multi-copy vector, we found that pMV261-*disA* caused inferior survival compared to the empty plasmid strain, further highlighting the role of c-di-AMP in terms of functional interception of RecA, which is a major protein involved in DNA damage repair. Conversely, it was also observed that, compared to *M. smegmatis* WT, the *M. smegmatis Δpde* strain (which has a high intracellular c-di-AMP level) showed fewfold more sensitivity to UV irradiation and the phenotype was reversed with complementation strain, highlighting the direct role of c-di-AMP on UV sensitivity (manuscript submitted). So, in the case of RadA, it was learned that there was no direct correlation between increased promoter induction and high copy number-driven better survival under UV irradiation; highlighting its indirect regulatory role in DNA repair. On the other hand, we hypothesize that DisA protein (as a scanning enzyme) could play an important role in DNA repair after UV irradiation-induced damage (Manikandan et al., 2018) when *disA* gene is preferably co-transcribed with *radA* using a stronger P2 promoter. Simultaneous expression of both RadA and DisA makes sure DisA can only work as a DNA scanning and repair enzyme, whereas the undesirable c-di-AMP synthesis is kept to a minimum level. Similarly, we checked ciprofloxacin sensitivity against *M. smegmatis* pMV261-*disA* and *M. smegmatis* pMV261-*radA* strain, and to our surprise, we found *M. smegmatis* pMV261-*radA* strain showed a significant level of sensitivity to 1 μg/ml. ciprofloxacin as measured by antibiotic disc diffusion assay **(Fig. 4b)**, unlike the *M. smegmatis* pMV261-*disA* strain and pMV261-*Empty* vector control strain. As we have seen very strong promoter induction of the P2 element with the same concentration of ciprofloxacin treatment, it is hypothesized to regulate c-di-AMP synthesis activity of the DisA enzyme to gain better fitness. In the case of ionic stresses of 500 mM NaCl and 500 mM KCl, pMV261-*disA* strain did not show better survival **(Fig. S4d)** compared to the blank vector control which is factually related to the unaltered expression of *disA* core promoter under the same conditions **(Fig. S4b)**. But, when we checked the same salt tolerance phenotype with *M. smegmatis* pMV261-*pde* strain, the survival percentage improved significantly **(Fig. S4d)**.

**Figure 4.**
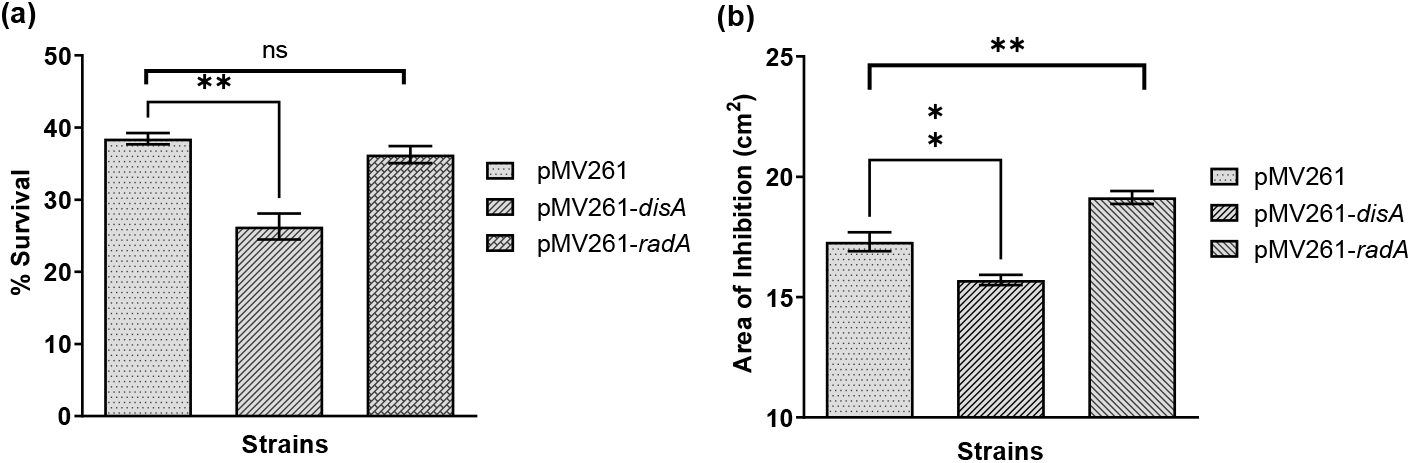
Stress survival assays **(a)** Percentage survival of mentioned strains was calculated after 0.15 mJ/cm^2^ UV irradiation. **(b)** ciprofloxacin sensitivity of the same strains is estimated by antibiotic disc diffusion assay. The graphs were plotted using GraphPad Prism 9. *** = P < 0.001; ** = P < 0.01; * = P < 0.05.

### High c-di-AMP and ppGpp regulate *disA* and *pde* core promoters

From earlier studies, it was known that high c-di-AMP concentration is toxic for cells (Tang et al., 2015) and hence intracellular c-di-AMP concentration fine-tuning is critical for the normal growth of the bacteria. Given this, we wanted to check if there is any transcriptional regulation of *disA* promoter that happens involving the c-di-AMP molecule itself (autoregulation). Steady-state homeostasis of c-di-AMP can also be achieved by regulating the promoter of the *pde*, the protein responsible for the degradation of c-di-AMP. We constructed a similar GFP-based transcriptional fusion construct by taking 300 bp upstream of *pde* (*P*_*pde*_) start codon and fused to GFP and confirmed steady state expression at different growth phases. Next, promoter induction of both *disA* and *pde* core promoters was checked in different strain backgrounds having low, high, or no c-di-AMP. It was observed that *disA* core promoter induction got altered in presence of high c-di-AMP concentration; precisely, our data suggested that *disA* promoter induction got significantly inhibited in *Δpde* strain in the late exponential phase **(Fig. 5a)**. This observation could be justified by autoregulation of c-di-AMP on *disA* core promoter after reaching a certain threshold concentration of intracellular c-di-AMP, beyond that it could be detrimental for cells. Similarly, we checked the *pde* core promoter induction at different strain backgrounds with low, normal, and high c-di-AMP concentrations and found out that there is no significant deviation of GFP intensity across all background strains **(Fig. 5b)**.To gain some insights into any possible crosstalk mechanism (at the transcription level) through an unidentified cascade between two second messengers c-di-AMP and (p)ppGpp in *M. smegmatis*, we checked the expression profile of *disA* and *pde* core promoter in (p)ppGpp overexpressing strain (*M*.*smegmatis+* pMV261*-rel*) (Petchiappan et al., 2020). Our data suggested that with the overexpression of the Rel enzyme, both the *disA* and *pde* genes’ core promoter induction remains largely unaltered compared to the corresponding blank vector controls **(Fig. 5c)**, which indirectly suggests increased (p)ppGpp level inside the cells does not contribute to any promoter level regulation of c-di-AMP synthesis and degradation.

**Figure 5.**
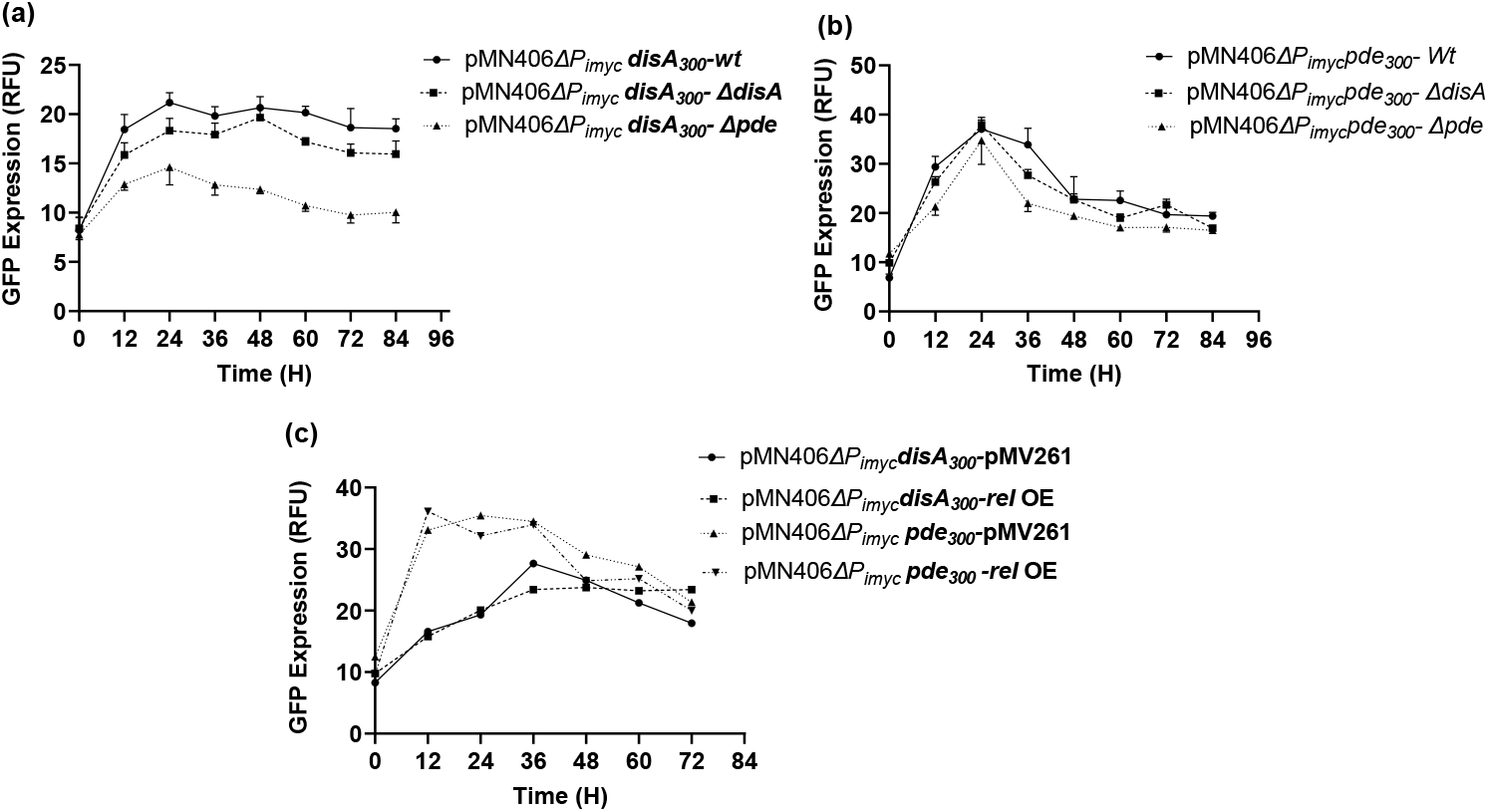
Promoter activity of P_*disA*300bp_ **(a)** P_*pde*300bp_ **(b)** is shown in different strain backgrounds having low, high, or no c-di-AMP. **(c)** Promoter activity of P_*disA*300 bp_ and P_*pde* 300 bp_ is measured in presence of high intracellular (p)ppGpp concentrations. Error bars are shown to validate results across biological replicates.

## Discussion

Bacterial polycistronic operons (Okuda et al., 2007; Zhao et al., 2019) can be described as a cluster of genes (often with a related function) that can be transcribed together to form a single stretch of mRNA and then translate into multiple proteins. Our group was particularly interested to study the regulation of c-di-AMP synthesis *M. smegmatis*, which is governed by a single synthetase enzyme DisA. The gene *disA* was shown before to exist in an operon where the other two genes are *radA* and *lpqE*. There were three major considerations when we initiated this study to understand the so-called sub-operon arrangement in *lpqE-radA-disA* operon in *M. smegmatis*. First, the presence of several transcription start sites (TSS), sigma factor consensus sequences(Junier et al., 2016; Yu et al., 2018) and distinct RBS within intergenic regions of genes could indicate the possibility of independent expression of the genes in the operon; second, the enzyme DisA being a dual functional protein (Fahmi et al., 2017) involved in both c-di-AMP synthesis as well as DNA damage monitoring, so was there any stress-induced transcriptional regulation of *radA* gene to shut off one activity (c-di-AMP synthesis) and keep the other (DNA binding and scanning) of DisA intact at the post-translational level (RadA-DisA interaction); third, to find out the specific conditions to justify the biological significance relating the (possibility of)- and (need for)-such gene-specific individual promoter induction within an operon.

As the precise control of c-di-AMP level is a prerequisite for the normal growth of cells (Fahmi et al., 2017) and DisA enzyme having dual functions, the expression and/or c-di-AMP synthesis activity of this multi-domain enzyme needs to be well coordinated. In this study, we have identified a sub-operon arrangement of 3 gens *lpqE, radA* and *disA*, based on distinct gene-specific sigma factor recognition sites, transcription start sites, and RBS. The presence of 3 different putative *sigB* recognition sequences upstream of the *disA* gene (within *radA* ORF) could be significant under stress conditions when un-inhibited c-di-AMP synthesis is required using the *disA* gene’s native TSS with a short 5’-UTR having a minor regulatory role **(Fig. 1b)**. On the other hand, when c-di-AMP synthesis is required at the basal level, using alternative housekeeping sigma factors, RNAP might prefer to use any upstream TSS sites which will have a long 5’-UTR region of mRNA facilitating the formation of a potential stem-loop structure **(Fig. S5)**, resulting in the minimal translational activity of *disA*. This observation further highlights the importance of the independent nature of the expression of the *disA* gene from the rest of the operon. Subsequently we validated the promoter activity of three putative promoter elements during normal growth to validate our bioinformatic observation. After that, it was important to check the comparative activity of the core and extended promoter of the *disA* gene under non-optimized growth conditions and we found a definite induction of the core promoter under carbon limitation conditions. Further, we were interested to identify stress-specific induction of the individual core promoters and tried to link the induction pattern with tolerance phenotype using DisA and RadA overexpression strains. We also explored if there was any *in vivo* auto-regulation involved with core *disA* and core *pde* promoter and we found that *P*_*disA*_ is transcriptionally downregulated in presence of high c-di-AMP inside cells, whereas no such regulation was observed in *P*_*pde*_ with variable intracellular c-di-AMP concentrations. We further measured the same promoter’s activity to know if c-di-AMP homeostasis gets disturbed in presence of an elevated concentration of (p)ppGpp and found no significant change in promoter activity. Future studies will be aimed to confirm such regulatory and functional crosstalk phenomenon of different second messengers in *M. smegmatis*. Finally, *disA* ORF sequence analysis confirmed multiple methionine (ATG) which could act as alternative start codons in-frame with the stop codon of the gene **(Fig. S6a)**. We did a structural modeling analysis to find out how the overall folding structure would have been impacted by those truncated proteins. As some of the truncated proteins would theoretically have a very short chain of amino acids, we only considered the one which lacked a substantial portion of the DAC domain and had potential RBS and sigma factor recognition sites upstream. One of them is the DisA truncated protein without the full-length DAC domain, more specifically having a deletion of the first 83 amino acids of the full-length DisA protein. This particular truncated region of the gene was more interesting than the rest because of the presence of a near-perfect RBS upstream of the start codon and a putative *sigE* promoter recognition sequence **(Fig. S6a)**. To find out, if there is any possible promoter element driving transcription of the downstream ORF, we cloned the upstream region of the alternate *disA* ORF up to the original(full-length) start codon of *disA* gene (∼246bp) upstream of GFP in the same reporter plasmid (P5 construct) **(Fig. S6b)**. Consequently, we checked the GFP induction of this transcriptional fusion construct (P5) against different relevant stresses and found no significant induction except UV stress. With UV exposure (0.15 mJ/cm^2^), to our surprise, the promoter was induced ∼2 fold **(Fig. S6c)** pointing toward our hypothesis of alternate promoter and ORF arrangement within the full-length ORF of *disA*. The physiological relevance of the truncated protein needs to be confirmed in the future. All in all, our data identified and characterized the sub-operon promoter arrangement of *disA* in *M. smegmatis* to counter stresses and distinct mechanisms to regulate c-di-AMP levels *in vivo*.

## Abbreviations

GFP: (Green Fluorescent Protein)
RFU: (Relative Fluorescence Unit)
FU: (Fluorescence Unit)

## Acknowledgments

We thank Prof. Ajitkumar, IISc, Bangalore for providing the pMN406 plasmid which was the backbone for all transcriptional fusion constructs. We thank Prof. Dipankar Chatterji, IISc for valuable feedback on the work. We also acknowledge Mr. Phani R.K. Behra for the bioinformatics study in the manuscript. AG thanks the Ramalingaswami Re-entry Fellowship and the Department of Biotechnology (DBT), Government of India, for funding this work (BT/RLF/Re-entry/31/2017). MS, AP, and VC acknowledge the Department of Biotechnology (DBT), Government of India for their fellowships.

## Statements & Declarations

### Author contributions

MS and AG contributed to the conception and design of the study. MS, AP, VC and AG performed the experiments. MS, AP and AG participated in data analysis and interpretation. MS and AG wrote the manuscript.

### Ethics approval

NA

### Consent to participate

NA

### Consent to publish

NA

