## Supplemental figures and tables for "Sub-operon promoter arrangement of *disA* facilitates c-di-AMP homeostasis and selective stress responses in *M. smegmatis*"

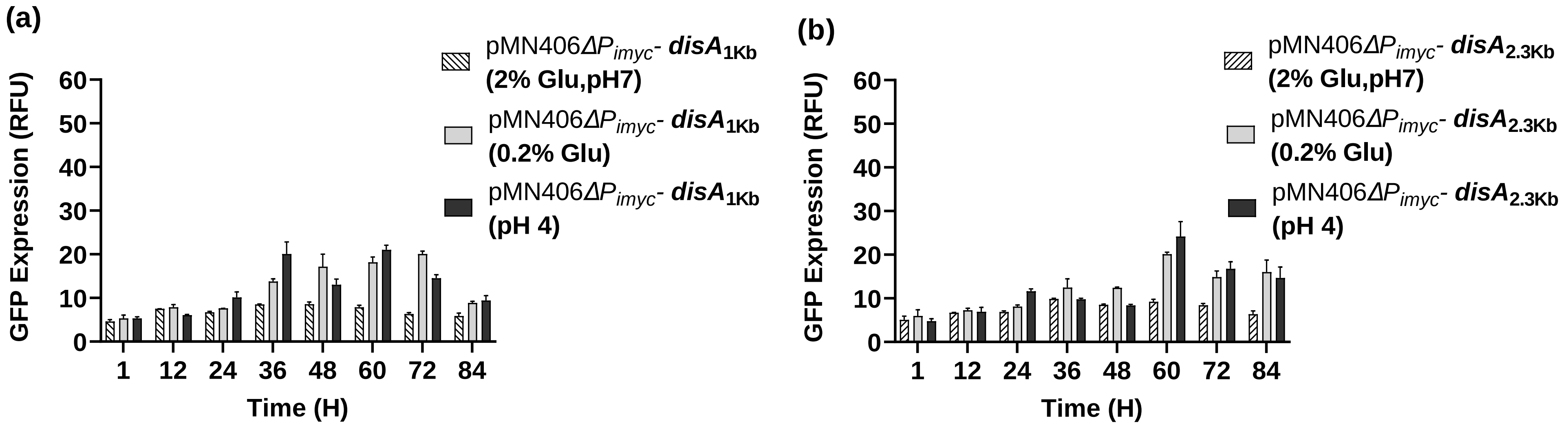


**Figure S1:** Promoter activity (a) *disA*1Kb and (b) *disA*2.3Kb fusion constructs are measured in a kinetic manner upon carbon stress and pH stress.

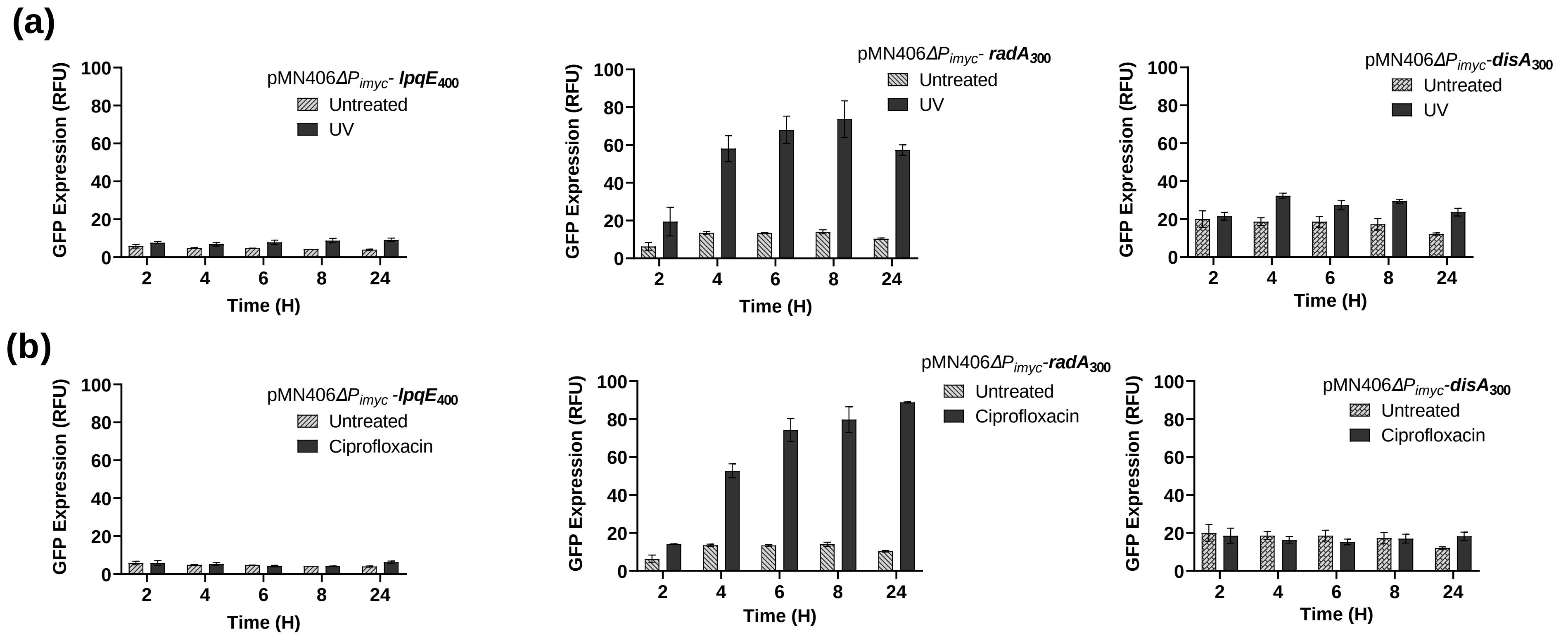


**Figure S2:** Individual promoter activity of P*lpqE* (left panel) , P*radA* (middle panel) and P*disA* (right panel) constructs are measured upon **(a)** 0.15 mJ/ cm2 UV irradiation and **(b)** 1µg/ml Ciprofloxacin stress, readings are taken in kinetic manner upto 24 hours. The graphs were plotted using GraphPad Prism 9. *** = P < 0.001; ** = P < 0.01; * = P < 0.05.

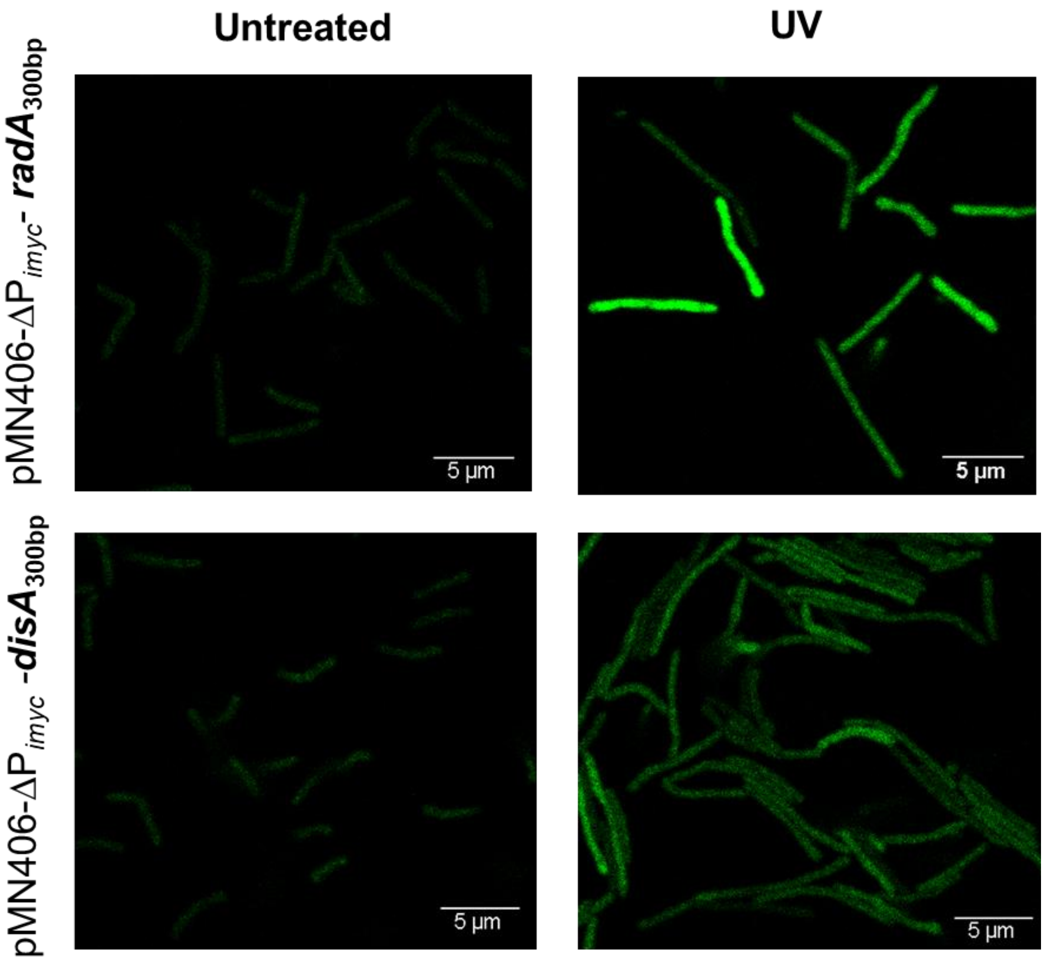


**Figure S3:** Representative microscopy images showing promoter induction of promoter activity of P*radA* and P*disA* before and after UV stress.

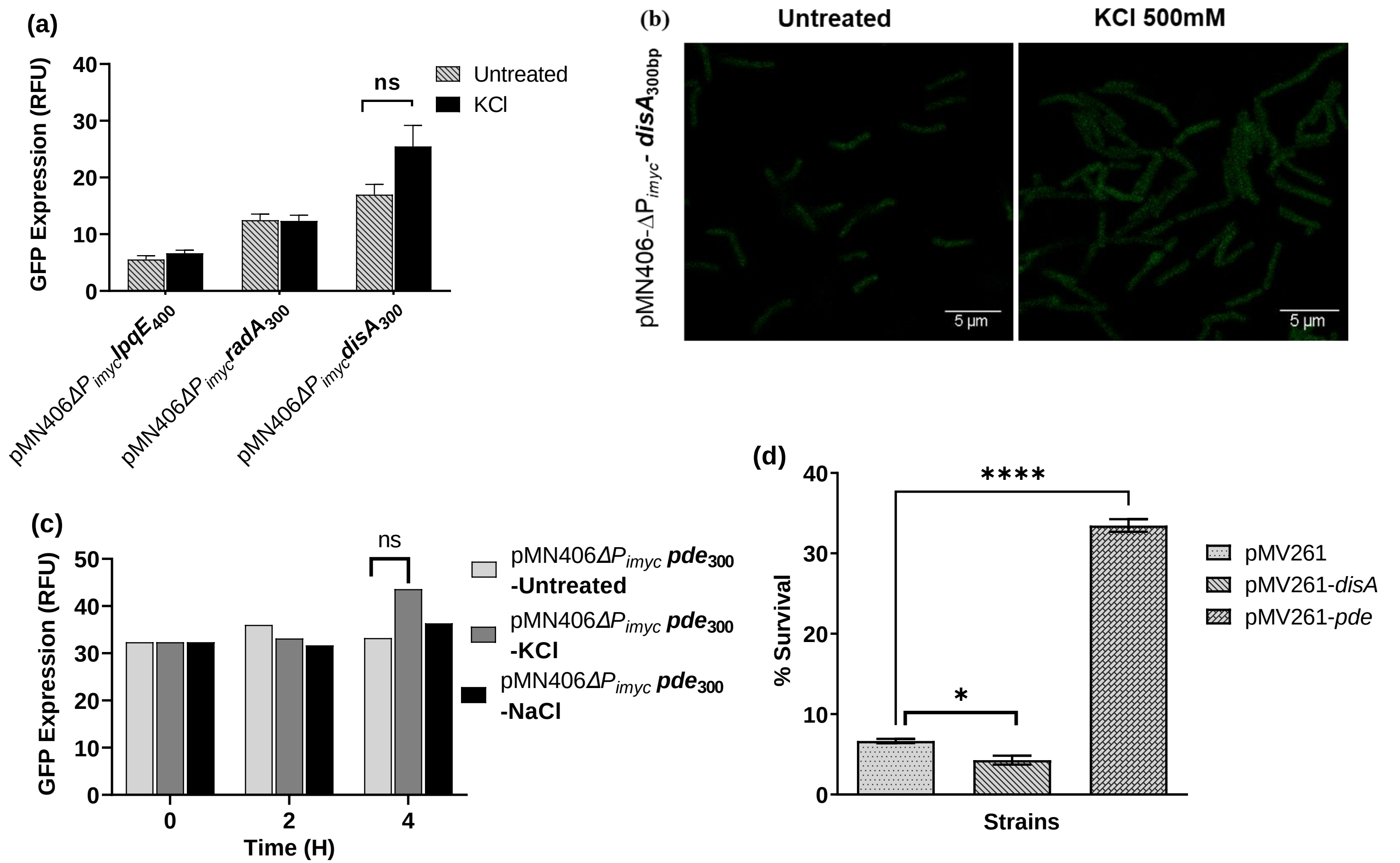


**Figure S4**: (A) Individual promoter activity of P*lpqE* (left panel) , P*radA* (middle panel) and P*disA* (right panel) constructs are measured upon salt stress, (B) Microscopy images showing hardly any promoter induction of P*disA* construct after salt stress. (C) Promoter activity of P*pde* after NaCl and KCl treatment shown no significant induction, (D) Percentage survival of mentioned strains is calculated after 500 mM NaCl treatment.

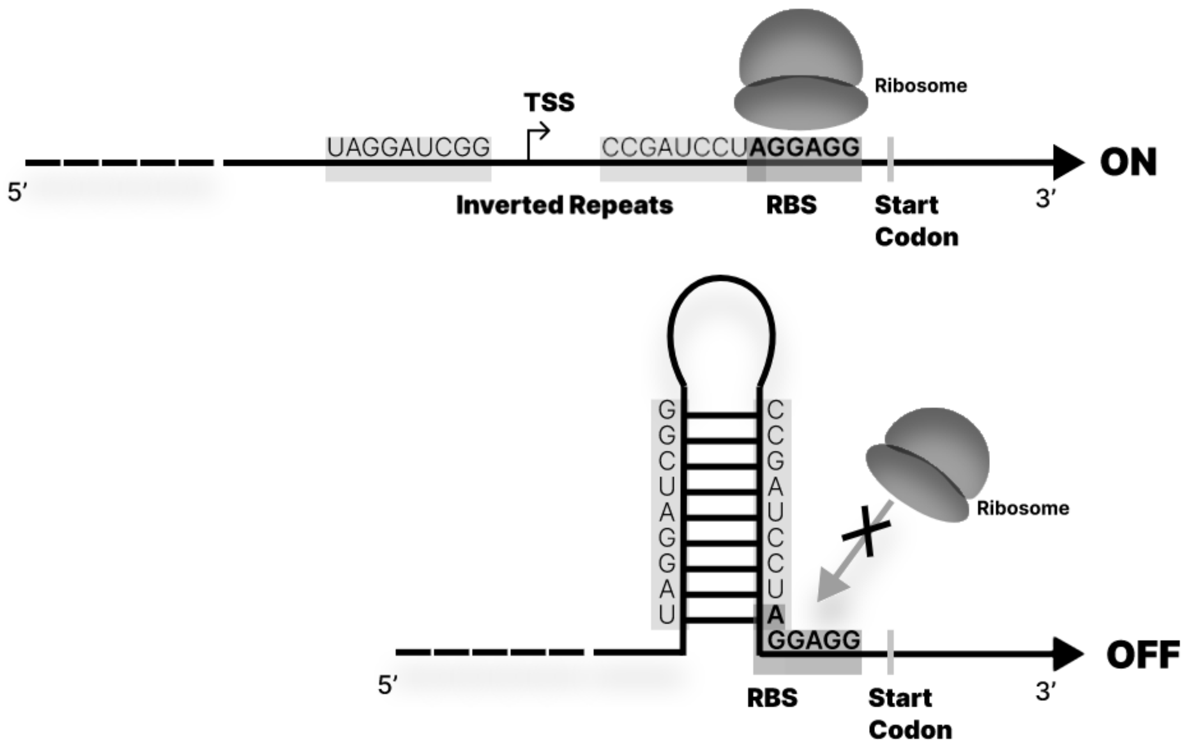


**Figure S5:** Schematic illustration of inverted repeats which is present upstream of *disA* gene and how it can contributes to hairpin structure formation resulting in translational inhibition.

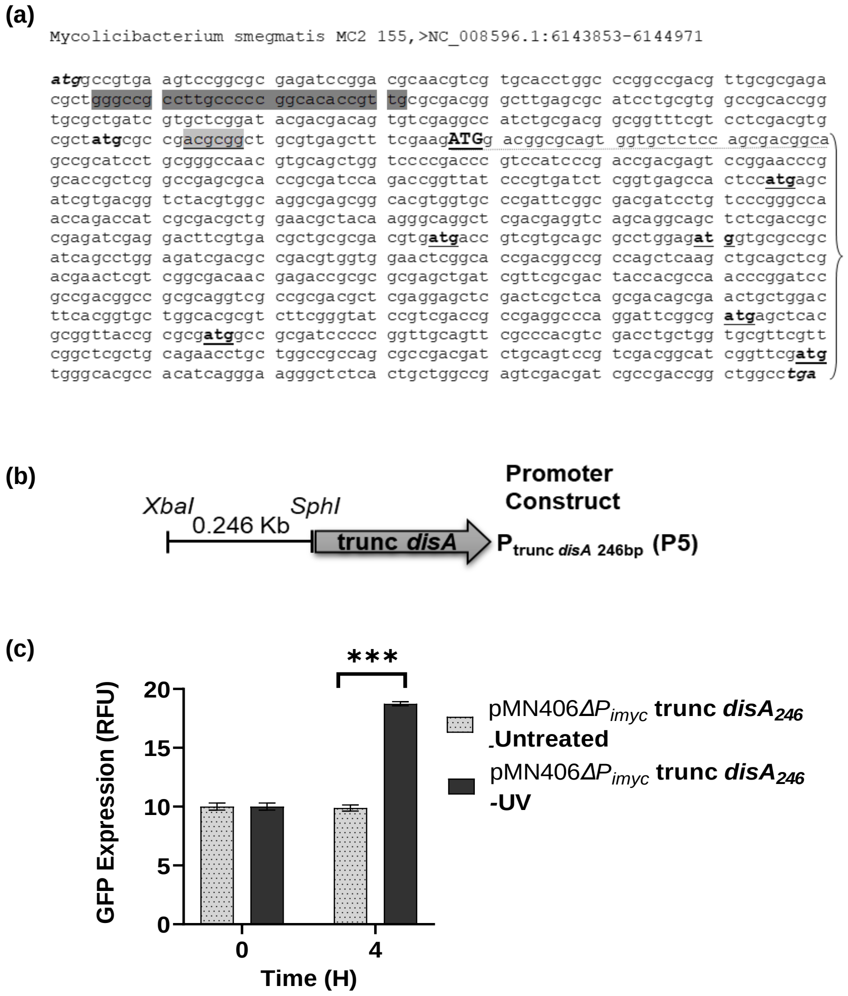


**Figure S6: (a)** DNA sequence of *disA* gene shows the presence of multiple "atg" codons (underlined) within the *disA* ORF. The start codon in capital letter (ATG) shows the start of truncated DisA protein, upstream sequence analysis highlights putative RBS and sigma factor recognition sequences (light grey and deep grey respectively), **(b)** schematic representation of truncated *disA* promoter fusion construct and **(c)** The promoter induction profile corresponding to the truncated *disA* region ($P_{disA\ trunc}$) is shown upon 0.15 mJ/ cm$^2$ UV irradiation.

**Table S1.** List of Bacterial Strains, plasmids and promoter fusion constructs used in the study

| Strains | Description | References |
|---|---|---|
| *M. smegmatis* Wild type | *M. smegmatis mc2155* | Laboratory stock [1] |
| *E. coli* DH5α | *E. coli* host strain for DNA cloning | Laboratory stock |
| *M. smegmatis ΔdisA* | *M. smegmatis* mc2 155 strain knock out for *disA*; KanR | Laboratory Stock |
| *M. smegmatis Δpde* | *M. smegmatis* mc2 155 strain knock out for *pde*; KanR | Laboratory Stock |
| ***Vectors:*** | | |
| pMN406-ΔPimyc | Mycobacterial shuttle vector pMN406 containing *mycgfp2+* gene, devoid of imyc promoter used for transcriptional fusion, HygR | Kind Gift from Prof. Ajitkumar, IISc Bengaluru. [2] |
| pMV261 | Mycobacterial vector used for cloning, KanR | Laboratory stock |
| pMV261-*disA* | pMV261 containing *disA* gene under the regulation of constitutive hsp promoter, KanR | Laboratory Stock |
| pMV261-*pde* | pMV261 containing *pde* gene under the regulation of constitutive hsp promoter, KanR | Laboratory Stock |
| pMV261-*radA* | pMV261 containing *radA* gene under the regulation of constitutive hsp promoter, KanR | This Study |
| pMV261-*rel* | pMV261 containing *rel* gene under the regulation of constitutive hsp promoter, KanR | Laboratory Stock |
| **Transcriptional fusion constructs in pMN406-ΔPimyc:** | | |
| pMN406-ΔPimyc- *disA*$_{300}$ | pMN406-ΔPimyc containing 314 bp *disA* upstream region, HygR | This Study |
| pMN406-ΔPimyc- *pde*$_{300}$ | pMN406-ΔPimyc containing 324 bp *pde* upstream region, HygR | This Study |
| pMN406-ΔPimyc- *disA*$_{1KB}$ | pMN406-ΔPimyc containing 1000bp *disA* upstream region, HygR | This Study |
| pMN406-ΔPimyc-*disA*$_{2.3KB}$ | pMN406-ΔPimyc containing 2337 bp *disA* upstream region, HygR(full length) | This Study |
| pMN406-ΔPimyc-*lpqe*$_{400}$ | pMN406-ΔPimyc containing 400 bp *lpqe* upstream region, HygR | This Study |
| pMN406-ΔPimyc-*radA*$_{300}$ | pMN406-ΔPimyc containing 300 bp *radA* upstream region, HygR | This Study |
| pMN406-ΔPimyc-*hsp60* | pMN406-ΔPimyc containing Hsp60 promoter region cloned using XbaI and PacI, HygR | Laboratory stock |
| pMN406-ΔPimyc-trunc *disA*$_{246}$ | pMN406-ΔPimyc containing first ~246 bp of *disA* gene, HygR | This study |
| pMN406ΔPimyc-*disA*$_{300}$-*ΔdisA* | pMN406ΔPimyc-*disA*$_{300}$ is transformed in *M. smegmatis ΔdisA*; HygR , KanR | This Study |
| pMN406ΔPimyc-*disA*$_{300}$-*Δpde* | pMN406ΔPimyc-*disA*$_{300}$ is transformed in *M. smegmatis Δpde*; HygR , KanR | This Study |
| pMN406ΔPimyc-*disA*$_{300}$- pMV261 *rel* | pMN406ΔPimyc-*disA*$_{300}$ is co-transformed with pMV261 *rel* in *M. smegmatis* Wild type; HygR ,KanR | This Study |
| pMN406ΔPimyc-*pde*$_{300}$-*ΔdisA* | pMN406ΔPimyc-*pde*$_{300}$ is transformed in *M. smegmatis ΔdisA*; HygR ,KanR | This Study |
| pMN406ΔPimyc-*pde*$_{300}$-*Δpde* | pMN406ΔPimyc-*pde*$_{300}$ is transformed in *M. smegmatis Δpde*; HygR , KanR | This Study |
| pMN406ΔPimyc-*pde*$_{300}$- pMV261 *rel* | pMN406ΔPimyc-*pde*$_{300}$ is co-transformed with pMV261 *rel* in *M. smegmatis* Wild type; HygR , KanR | This Study |

**Table S2.** List of oligonucleotides used in this study. Restriction enzyme sites, wherever present, are shown in bold letters.

| Primer name | Oligo Sequence (5’-3’) | Use in the study |
|---|---|---|
| *disA*_Psd5b_300F | CGAT**TCTAGA**GCAGACCTCGCGGTG | Cloning/Sequencing |
| *disA*_pSD5bR | ATCG**GCATGC**CAGCCCTCCTAGGAT | Cloning |
| *disA*_1000bp_XbaI_F | ATGC**TCTAGA**CGCACGATCAGGTGTACCTG | Cloning |
| *disA*_1000BP_XbaI_R | TAGC**TCTAGA**CAGCCCTCCTAGGATCGGC | Cloning |
| *radA* UTR F | ATAT**TCTAGAAAGCTT**TCTGATTCGAGCCCCTTTCG | Cloning/Sequencing |
| lpqE_400us_XbaI_FW | CGAT**TCTAGA**ACCTTGAGGACGAGATATTC | Cloning |
| lpqE_400us_SphI_REV | ATCG**GCATGC**TCAAGCCTCCTGAACAGGTT | Cloning |
| *radA*_300us_XbaI_FW | CGAT**TCTAGA**CCGCCGACGGCGTGCTGATC | Cloning/Sequencing |
| *radA*_300us_SphI_REV | ATCG**GCATGC**ACGCCGTGACAGTATCGGCA | Cloning |
| mspde_pSD5b_300 F | CTAG**TCTAGA**GTACTACACCGTGCT | Cloning |
| mspde_pSD5bR | TAGC**GCATGC**CCTCAGCGTTCGTCC | Cloning |
| Gfp_SphI_fw: | CTAG**GCATGC**ATTCGAAGGGCGAGGAGCTGTT | Cloning |
| Gfp_HindIII rev: | ATCG**AAGCTT**CTACTTGTACAGCTCGTCCATG | Cloning |
| *disA*_TRUNC_XbaI_F | ATGC**TCTAGA**ATGGCCGTGAAGTCCGGC | Cloning |
| *disA*_TRUNC_SphI_R | ATCG**GCATGC**CTTCGAAAGCTCACGCAG | Cloning |
| Pmv261_radAF_HindIII | ACTG**AAGCTT**AGGAGGCGGCGTGGCCGGTTCG | Cloning |
| Pmv261_radAR_HindIII | ACTG**AAGCTT**CTATTGGGCACCCGCA | Cloning/Sequencing |
| GFPM2+ term rev | ACTTGTGGCCGTTGACGTC | Sequencing |
| *disA*_550bp_US_Rev | CGCGGTACCGGGCACCGCGA | Sequencing |
| pMN406 seq FW | GGCCGATTCATTAATGCAGC | Sequencing |
| pMV261_seq_R | GATGATATATTTTTATCTTGTGC | Sequencing |
